# OligoN-design: A simple and versatile tool to design specific probes and primers from large heterogeneous datasets

**DOI:** 10.1101/2025.11.04.685038

**Authors:** Miguel M. Sandin, Marie Walde, Nicolas Henry, Irene Forn, Nathalie Simon, Cédric Berney, Ramon Massana, Daniel J. Richter

## Abstract

High-throughput environmental DNA sequencing has ushered ecological and evolutionary studies into the big data era. With thousands to millions of DNA sequences, designing taxon-specific oligonucleotides is a current bottleneck of molecular studies that rely on primers for Polymerase Chain Reactions (PCRs) or probes for Fluorescence *in situ* Hybridization (FISH). No software currently exists to design specific oligonucleotides starting from a custom set of sequences. Existing tools rely on specific databases, alignments or phylogenetic trees, or cannot accommodate increasingly large molecular environmental datasets. Here we present oligoN-design, a versatile tool to design oligonucleotides specific to a set of target sequences while minimizing predicted binding to non-target sequences. OligoN-design is simple, reproducible, and adaptable to high-throughput sequencing data analyses. It requires only two fasta files as input, one containing target taxa and the other containing non-target taxa. Using standard bioinformatic formats, it integrates easily with other tools such as BLAST, VSEARCH or MAFFT. OligoN-design allows a range of strategies that we present in detail, from an unsupervised end-to-end usage all the way to a detailed and thorough expert usage. Starting with large, comprehensive ribosomal databases that are widely used by the community (i.e., PR2, SILVA) and the unsupervised function, we were able to replicate known taxa-specific oligonucleotides in under 30 minutes and up to 6 Gb of RAM on a personal laptop. OligoN-design v1, available at github.com/MiguelMSandin/oligoN-design under GNU General Public License version 3.0, is easily installed via bioconda bioconda.github.io/recipes/oligon-design/README.html.

## 1. Introduction

Environmental DNA sequencing has unveiled an enormous amount of previously unseen diversity, from uncultured taxa within known lineages to a panoply of novel groups at different taxonomic scales, from prokaryotes to eukaryotes. Environmental sequence datasets produced today include short hypervariable regions from the universal Small SubUnit of the ribosomal DNA (SSU rDNA: 16S rDNA in prokaryotes and 18S rDNA in eukaryotes; e.g., Pernice et al., 2015; Alberti et al., 2017), but also full length rDNA from metagenomes and metatranscriptomes (e.g., Krabberød et al., 2025), or direct long-read sequencing (e.g., Jamy et al., 2019). The combination of different approaches and increasingly large environmental datasets has enabled a more detailed characterization of both novel environmental groups and well-defined taxonomic groups. Many of these groups can only be studied through culture-independent molecular approaches (e.g., Massana et al., 2014; Sandin et al., 2025), and many others escape universal primers (Vaulot et al., 2022). The design of specific oligonucleotide primers or probes (short nucleic acid sequences of ∼16-20 bases complementary to RNA or DNA signature regions of the target) could therefore accelerate the study of their biodiversity and biogeography.

Given the genetic heterogeneity of environmental datasets, and the large number of sequences they can include (from 100s to 100,000s), oligonucleotide design is a current bottleneck for molecular-based ecological and evolutionary analysis of specific lineages. Both the Polymerase Chain Reaction (PCR) and Fluorescence in situ Hybridization (FISH) rely on specific complementary oligonucleotides to detect sequences of interest, and despite the fact that they are routine approaches, oligonucleotide design still requires manual and tedious work. Several software packages have been developed to aid in the design of specific oligonucleotides, such as ARB (Ludwig et al., 2004), primer3 (Untergasser et al., 2012), DECIPHER (Wright, 2016), PrimerMiner (Elbrecht and Leese, 2017), oli2go (Hendling et al., 2018) or OligoMiner (Beliveau et al., 2018), among others (Hendling and Barišić, 2019). However, these tools require at least one input that is not easily available for large environmental datasets: sequence alignments, phylogenetic trees, specific databases and/or genomes. In addition, their learning curves might be steep, with many different parameters that might be cryptic for the non-expert, they may allow for only a single input target sequence, they may poorly accommodate increasingly large and genetically heterogeneous molecular environmental datasets, or they are no longer available.

To overcome these limitations, we introduce oligoN-design, a simple open-source tool to design specific oligonucleotides to be used as primers for PCR, probes for FISH, or any other application. OligoN-design is simple, versatile and reproducible since the code and environment can be saved and rerun, accommodates large environmental, high-throughput and heterogeneous datasets, and does not require phylogenetic trees, alignments or specific databases as input. It uses common formats in bioinformatics, such as FASTA files and tab-delimited tables, so it can also be directly integrated with different tools such as VSEARCH (Rognes et al., 2016), BLAST (Camacho et al., 2009) or MAFFT (Katoh and Standley, 2013), among others. OligoN-design only requires two fasta files as input, one containing the sequences of the target set of taxa, and one containing the sequences of the excluding groups (i.e., all non-target sequences). OligoN-design is sufficiently versatile to be accessible to users of different bioinformatic expertise: it allows the novice user to run an unsupervised oligonucleotide search, while at the same time, an expert user might take advantage of advanced functions to inspect homology in specific regions of the target sequence. Earlier versions of the oligoN-design tool have been successfully used for designing both FISH probes (e.g., Sørensen et al., 2023) and PCR primers (e.g., Pardasani et al., 2025), demonstrating its practical use. Here, we present the rationale and structure of oligoN-design v1, show typical use cases and discuss its advantages and limitations.

## 2. The oligoN-design Tool

OligoN-design v1 is a collection of functions to ease the tedious work of oligonucleotide design. Probe and primer oligonucleotides are typically designed to be as specific as possible to the group of interest (the target) and potential pitfalls in their design include false-positive matches to non-target groups (e.g., Vaulot et al., 2022), low specificity of the oligonucleotide in suboptimal annealing conditions (e.g., Dieffenbach et al., 1993), self-binding of the oligonucleotide (hairpins, self-dimers; e.g., Nazarenko et al., 2002) or poor accessibility of the targeted region within the secondary structure of the targeted molecule (e.g., Behrens et al., 2003), among others (see e.g., Hendling and Barišić, 2019 for a review on in silico challenges and approaches).

To mitigate all of these issues, oligoN-design provides a series of individual steps linked together in a workflow. First, oligoN-design takes a **target** fasta file containing target sequences and searches for regions conserved in these sequences but absent in the non-target sequences contained in an **excluding** fasta file (Fig. 1). Once specific regions have been found, these can be further studied in order to predict mismatches, hairpins, self-dimers and/or the region accessibility based on the secondary structure, among other parameters. They can also be directly inspected against homologous regions within the **excluding** file for a more detailed and thorough examination of the position of the mismatches and their identity. Altogether, criteria based on these parameters are used to select candidate oligonucleotides for further empirical tests in the laboratory.

**Figure 1.**
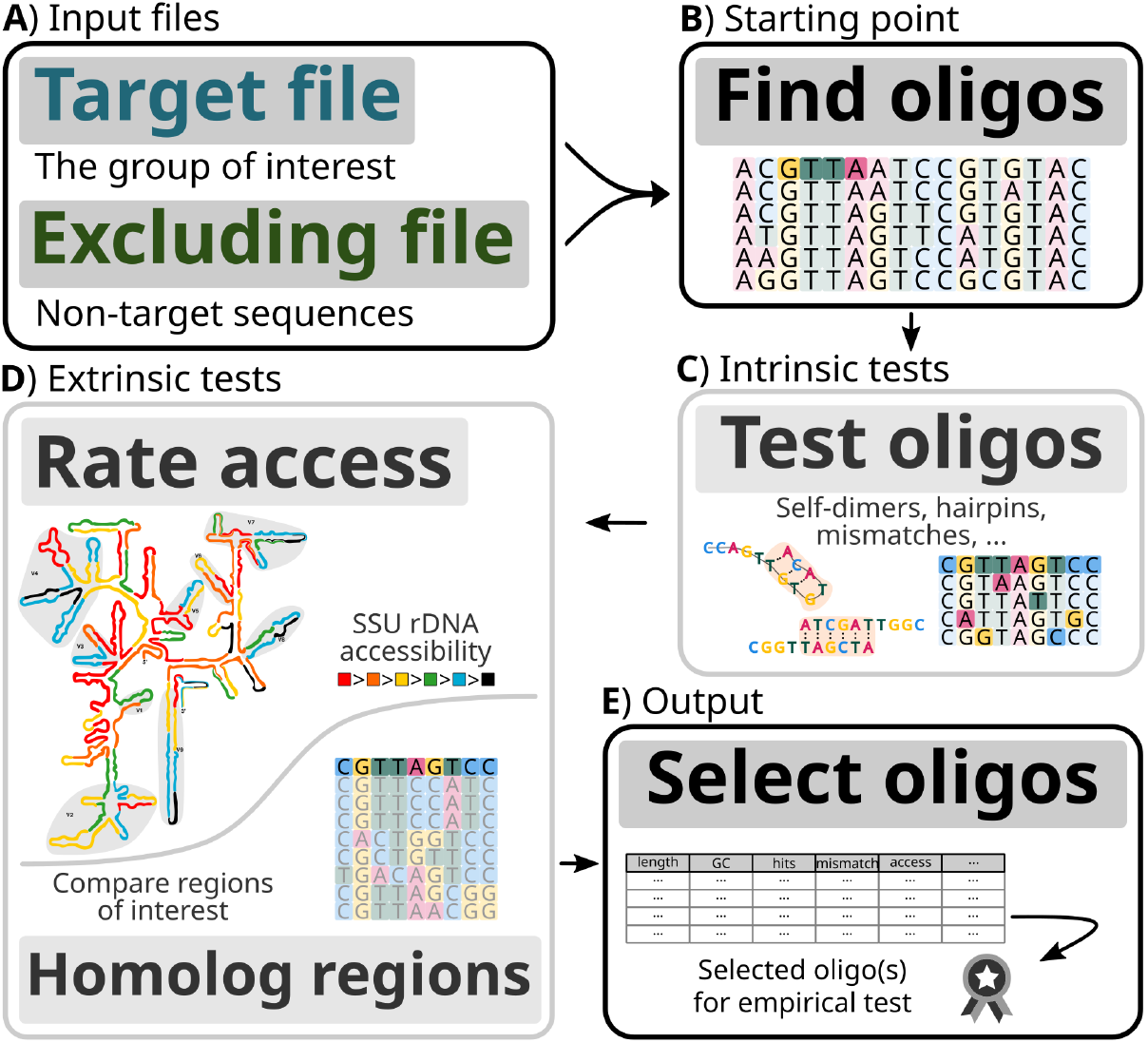
Schematic representation of the oligoN-design tool and workflow. Black boxes represent mandatory steps, gray boxes represent optional steps. Briefly, (**A**) from two input files containing the target sequences and the non-target sequences, (**B**) the oligoN-design tool starts by finding all potential oligonucleotides. These oligonucleotides can then be tested for (**C**) intrinsic properties such as self-dimers, hairpins or mismatches against the non-target sequences, and/or (**D**) extrinsic properties such as the accessibility in the gene’s secondary structure or compared against homologous regions in the non-target sequences. Lastly, (**E**) the best suitable oligonucleotides are listed based on all combined properties for further empirical test in the laboratory. For more details, please see the text and/or Figure S1.

Below is a brief description of the logic of oligoN-design, its main functions and how each function can be integrated to facilitate the design of specific oligonucleotides. For a detailed and complete description of all functions and fully documented replicable workflows please refer to the documentation manual, and for a graphical detailed overview of the tool see Figure S1. The oligoN-design v1 tool is mostly written in Python 3, relying on dependencies that are commonly installed in bioinformatics compute clusters, such as MAFFT (Katoh and Standley, 2013) for aligning sequences, HMMER (hmmer.org/) to search homologous regions and agrep (Wu and Manber, 1992) for approximate matching, or common python libraries such as Biopython (Cock et al., 2009), pandas (The pandas development team, 2024) and Matplotlib (Hunter, 2007). The tool is available at github.com/MiguelMSandin/oligoN-design under the GNU General Public License version 3.0 and can be easily installed from Bioconda (bioconda.github.io/recipes/oligon-design/README.html) together with all its dependencies.

### 2.1. A collection of versatile functions

Designing oligonucleotides is a diverse task that is approached from many different angles depending on the user’s experience. A novice user with limited experience in bioinformatic analysis and oligonucleotide design might simply be interested in the top 4 best-scoring oligonucleotides. However, a more experienced user might be interested in recalibrating search parameters, interacting with intermediate files, or applying different approaches or functions from other packages. The thorough user might even be interested not only in the number of mismatches, but in the position and the identity of the mismatched hit. And an expert user might be interested in inspecting the alignment manually. However given the sizes of environmental datasets, the visual inspection of an alignment could be challenging if not meaningless. In that case, the expert user might be interested in reducing the dataset to only the relevant homologous region. Altogether, from an unsupervised run to an expert usage, oligoN-design offers different functions to accommodate the user’s preferred workflow.

Here we suggest 4 different workflows (Fig. 2) to introduce the main functions and to showcase the versatility and integration of the oligoN-design v1 tool. These workflows go from an end-to-end unsupervised usage of the oligoN-design tool to an expert usage. A detailed and hands-on report of each workflow can be found in the documentation manual, and replicable example scripts can be found in the directory ‘pipelines’ from the github repository.

**Figure 2.**
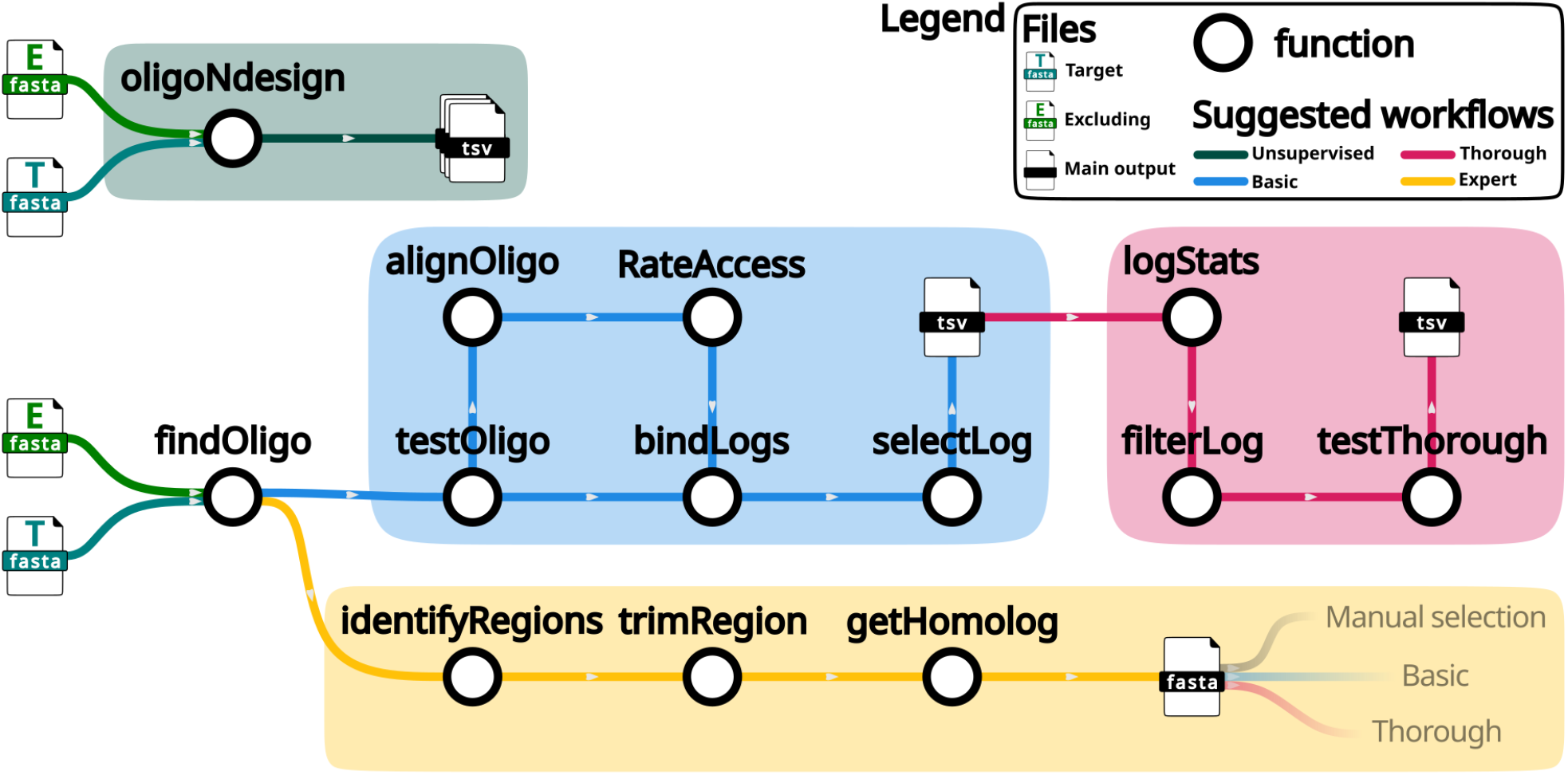
Different workflows suggested in this manuscript and detailed in the text. Briefly, the **unsupervised** workflow is the automatic implementation of the **basic** workflow, in which oligonucleotides are found and tested for mismatches and their accessibility is rated in the tertiary structure of the rDNA, to select the N (by default N=4) best scoring oligonucleotides. The **thorough** workflow starts from a tab delimited table (either resulting directly from the oligonucleotides found or from the final output of the basic workflow), filters only oligonucleotides of interest based on global statistics defined by the user, and applies a thorough test to explore the position and identity of the mismatches. Lastly, the **expert** workflow identifies potential regions of interest (as opposed to given oligonucleotides of defined lengths) and extracts homologous regions to later apply a manual selection of the oligonucleotides, or instead to use the output as a starting target and excluding files as input to a new run of the basic and/or thorough workflow.

### 2.2. Unsupervised run

The simplest usage of the oligoN-design tool is running the unsupervised wrapper function **oligoNdesign** (Fig. 2, dark green workflow; corresponding to the functions of the basic workflow described below, with default options). Briefly, this function will search specific oligonucleotides to the **target** fasta file that are not in the **excluding** fasta file, test them, rate their accessibility in the SSU (if applicable), and select the 4 best scoring oligonucleotides. For practical examples of this workflow, see “4. Recommendations based on a qualitative performance” further down.

### 2.3. Basic workflow

The basic workflow is the supervised version of the **oligoNdesign** function. Here, the user can interact with all intermediate files, arguments and options, which allows tuning and optimizing user-defined thresholds according to the data and needs. The main starting point is the function **findOligo** (Fig. 2, blue workflow). By default it searches for specific regions of length 18 and 20 bases that are present in at least 80% of the sequences from the **target** file and in less than 1% of the sequences in the **excluding** file. These thresholds were chosen based on empirical evidence of the most common specific oligonucleotides properties but can be adapted by the user based on their experimental requirements (as for any other threshold within oligoN-design). As a result, a tab-delimited table is exported, containing the identified candidate oligonucleotides and several values such as the GC content and the number and proportion of hits in the **target** and **excluding** files (Fig. 3A).

**Figure 3.**
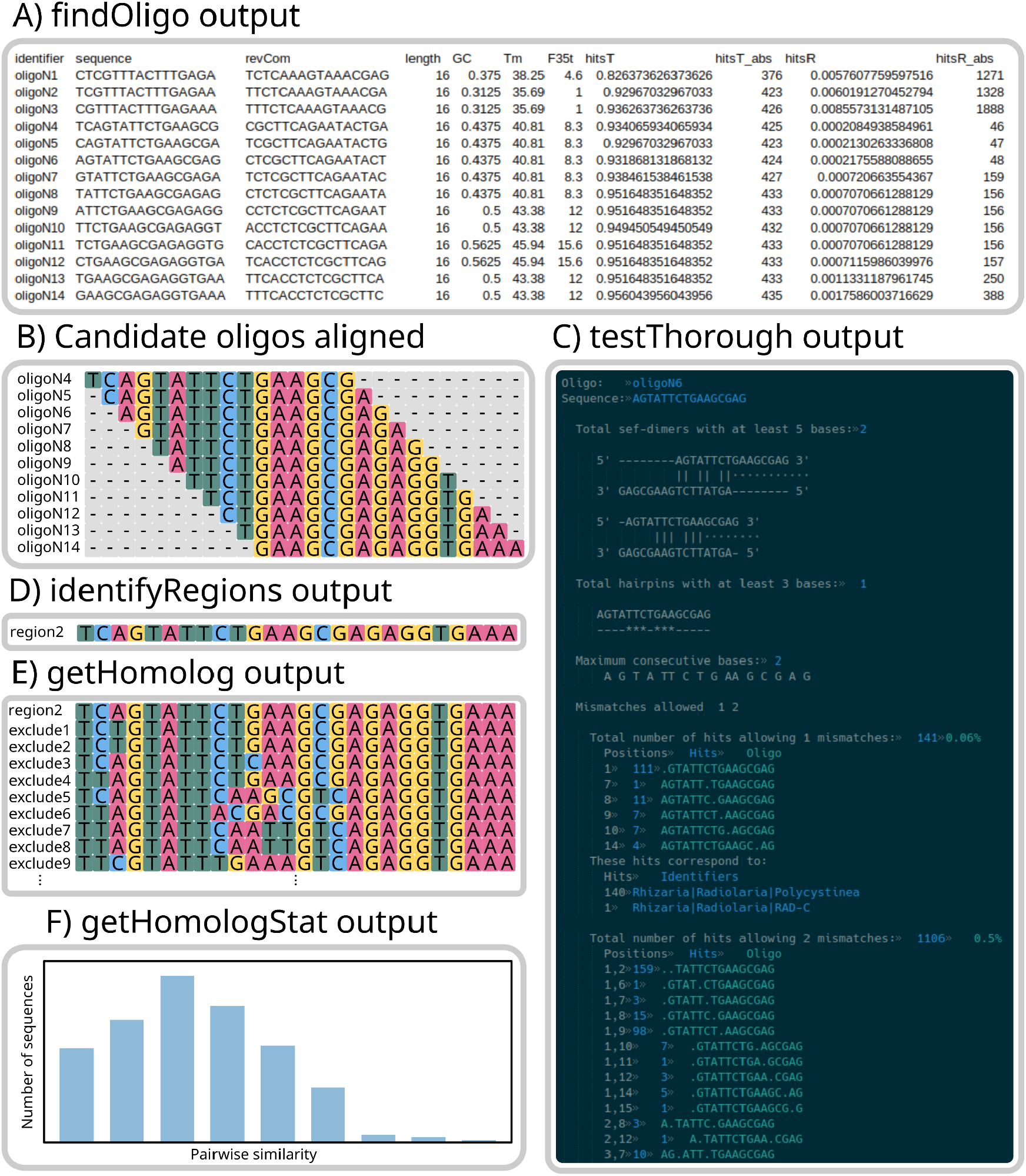
Example outputs of different functions of oligoN-design: **A**) a screenshot of a tab delimited table output from **findOligo**; **B**) a screenshot of the **testThorough** output; **C**) a schematic representation of the candidate oligonucleotides obtained from **findOligo** as sliding windows of length *k* from a region of length *>k*; **D**) a fasta file output from **identifyRegions**; **E**) a fasta file output from **getHomolog**; **F**) a plot prompted from **getHomologStats**.

The candidate oligonucleotides found with the **findOligo** function (Fig. 3B) are subsequently tested for mismatches to optimize specificity and minimize false-positive hybridizations, as follows: the function **testOligo** takes advantage of the ‘agrep’ program (Wu and Manber, 1992) for fast and approximate matching to test for mismatches allowing insertions and deletions against the **excluding** file. For datasets focusing on the SSU rDNA, candidate oligonucleotides can also be tested for their accessibility in the secondary structure of the SSU rDNA after the empirical results from (Behrens et al., 2003) with the function **rateAccess**. However, for other genes the user can also benefit from the function **rateAccess** if a custom accessibility table is provided (see documentation or help function for further details).

Selecting potential oligonucleotides from all candidate oligonucleotides will depend on the user’s preferences, experience, and most importantly, the question of the study. By default, all search functions from oligoN-design will output a tab-delimited table with values and parameters specific to the function used (see documentation manual for details about the specific output of each function). Different tables can be merged with the function **bindLogs**, and then the user can select the best oligonucleotides based on the resulting values (e.g., high specificity, high GC content, low hits to the **excluding** file allowing mismatches, highly accessible, etc). Additionally, the best scoring oligonucleotides can be ranked with the function **SelectLog**, and thus the novice user can select the best oligonucleotides without relying on subjective criteria.

### 2.4. Thorough design

More thorough tests can also be applied with the function **testThorough** (Fig. 2, magenta workflow, see Fig. 3C for an example output), including evaluating the risks of hairpin and homodimer formations, maximum consecutive bases, mismatches, their exact position in the oligonucleotide and the identity of the mismatched sequence. The desired position of mismatches depends on whether the oligonucleotide will be used as a hybridization probe (where central mismatches are favored) or as a PCR primer (where the most important mismatches are close to the 3’ end where DNA polymerization starts). Since these analyses are relatively computationally expensive, we suggest that the user filters the searched oligonucleotides and provides a small input file, for example providing the best scoring oligonucleotides from the basic workflow or from the **findOligo**’s output. This will yield a more efficient computation by only performing the thorough tests on oligonucleotides that will most likely be kept for empirical laboratory experiments. To do so, the user can explore the obtained parameters with the function **logStats**, that will produce summary statistics (i.e., percentiles, mean) from all different properties, and manually filter oligonucleotides of interest with the function **filterLog**.

### 2.5. Expert design

Lastly, and closest to the traditional approach to designing oligonucleotides, is the “simple” examination of the alignment. However, given the large number of available sequences, this approach is becoming increasingly impractical and a considerable amount of experience is needed. To reduce the size of the alignment to be examined, the oligoN-design tool can be used to identify specific regions of interest from the **target** file and search for homologous regions in the **excluding** file (Fig. 2, yellow workflow).

Candidate oligonucleotides are frequently sliding windows of a given size from specific regions (Fig. 3B) that can be grouped and identified with the **identifyRegions** function (Fig. 3D). Then, homologous regions from the **excluding** file are retrieved with the function **getHomolog** (Fig. 3E) using an HMM profile from the HMMER package (hmmer.org/). The resulting fasta file can be analyzed with **getHomologStats** to estimate the pairwise similarity of all sequences of a fasta file against the first sequence in the file to produce a histogram with the similarities (Fig. 3F). These regions can also be used as a starting point to apply a basic or a thorough workflow on highly curated **target** and **excluding** files, thus continuing with an automated and replicable pipeline. This approach speeds up computational analysis for testing mismatches, while at the same time allowing a greater variety of oligonucleotide lengths and visually inspecting homologous regions (e.g., checking for variable positions in the **target** file, mismatches against the **excluding** file, the position of the mismatches, etc).

### 2.6. Other complementary aspects and functions

Even though alignments are not strictly required for oligoN-design, we recommend that advanced users align, specifically, the **target** file to properly understand both the file and the group of interest (see “Good practices” on the documentation manual for further details). Such a practice will help the advanced user fine-tune parameters related to the specificity, and probably discard or focus on specific regions of the gene that have a low or a high coverage, respectively. However, large alignments are sometimes difficult to visualize; the function **alignmentConsensus** creates a consensus sequence to quickly see highly variable or conserved regions within the group of interest. And in the case that computing resources do not allow aligning a given file, the function **breakFasta** extracts all unique k-mers of length *k* and their presence abundance in the input file (including an option to plot a scatter plot with such abundances) to help the user select different oligonucleotide lengths. For further details about these and other auxiliary and helper functions, please see the “Quick overview of the different functions and their usage” on the documentation manual (page 6).

All these functions allow the user to assemble a fully reproducible and end-to-end workflow, while at the same time letting an expert user visually inspect regions of interest. This versatility allows the integration of other complementary and already available tools for oligonucleotide design, such as primer3 (Untergasser et al., 2012), a widely used programme that focuses on thermodynamic properties of the oligonucleotide.

## 3. Novelty and complementary aspects

OligoN-design complements existing software and helps the user streamline the process of oligonucleotide design in the age of environmental omics. OligoN-design focuses mostly on specificity, accommodating large and heterogeneous environmental datasets.

Of existing oligonucleotide design software, probably the most widely used for probe and primer design in ecological and evolutionary studies is the ARB project (Ludwig et al., 2004) integrated with the SILVA database of the SSU rDNA (Quast et al., 2013). Briefly, this interactive tool allows the user to navigate a phylogenetic tree (and a very large alignment) and to select the taxa of interest in the tree. Signature regions in the target sequences and mismatches in the non-target sequences can be visualized in alignments as well as in models of secondary gene structure. Although ARB has proven very successful for oligonucleotide design, it has a steep learning curve and many dependencies, and requires a previous alignment and a phylogenetic tree, making it difficult to accommodate custom databases. Primer3 (Untergasser et al., 2012), another widely used tool for oligonucleotide design (mostly in biomedical studies), incorporates accurate thermodynamic models to improve the prediction of the melting temperature and therefore minimize empirical optimization in the laboratory. However, primer3 only accepts one input sequence. Lastly, the R packages DECIPHER (Wright et al., 2014; Wright, 2016) and PrimerMiner (Elbrecht and Leese, 2017) allow designing target-specific oligonucleotides from alignments and creating databases in compressed native formats, yet permit a relatively low heterogeneity within the target sequences. In this landscape of oligonucleotide design software, the oligoN-design tool introduces a simple and versatile command-line approach designed to accommodate large, heterogeneous and custom environmental datasets into bioinformatic analysis.

## 4. Recommendations based on a qualitative performance

We checked the qualitative performance of oligoN-design by applying *a posteriori* search for commonly used oligonucleotides on large and comprehensive empirical databases including the EukRibo (v2; Berney et al., 2022), SILVA (v138.2.NR99; Quast et al., 2013) and the Protist Ribosomal Reference (PR2 v5.0.0; Guillou et al., 2013) databases of SSU sequences. More precisely, we searched for randomly-selected general and specific FISH probes described as “very good” in the Supplementary Table 1 from (Piwosz et al., 2021), using the unsupervised function **oligoNdesign**. The different scope of the databases yielded very different results when retrieving and identifying already known oligonucleotides (or FISH probes in this example; Table 1). While all three databases are meant to taxonomically assign reads from High Throughput Sequencing, they all differ in, for example, their quality controls or naming strategies, and therefore the input files must be independently curated. Below, a brief description of the yielded probes, the curation strategies, and further recommendations to the user.

**Table 1.**
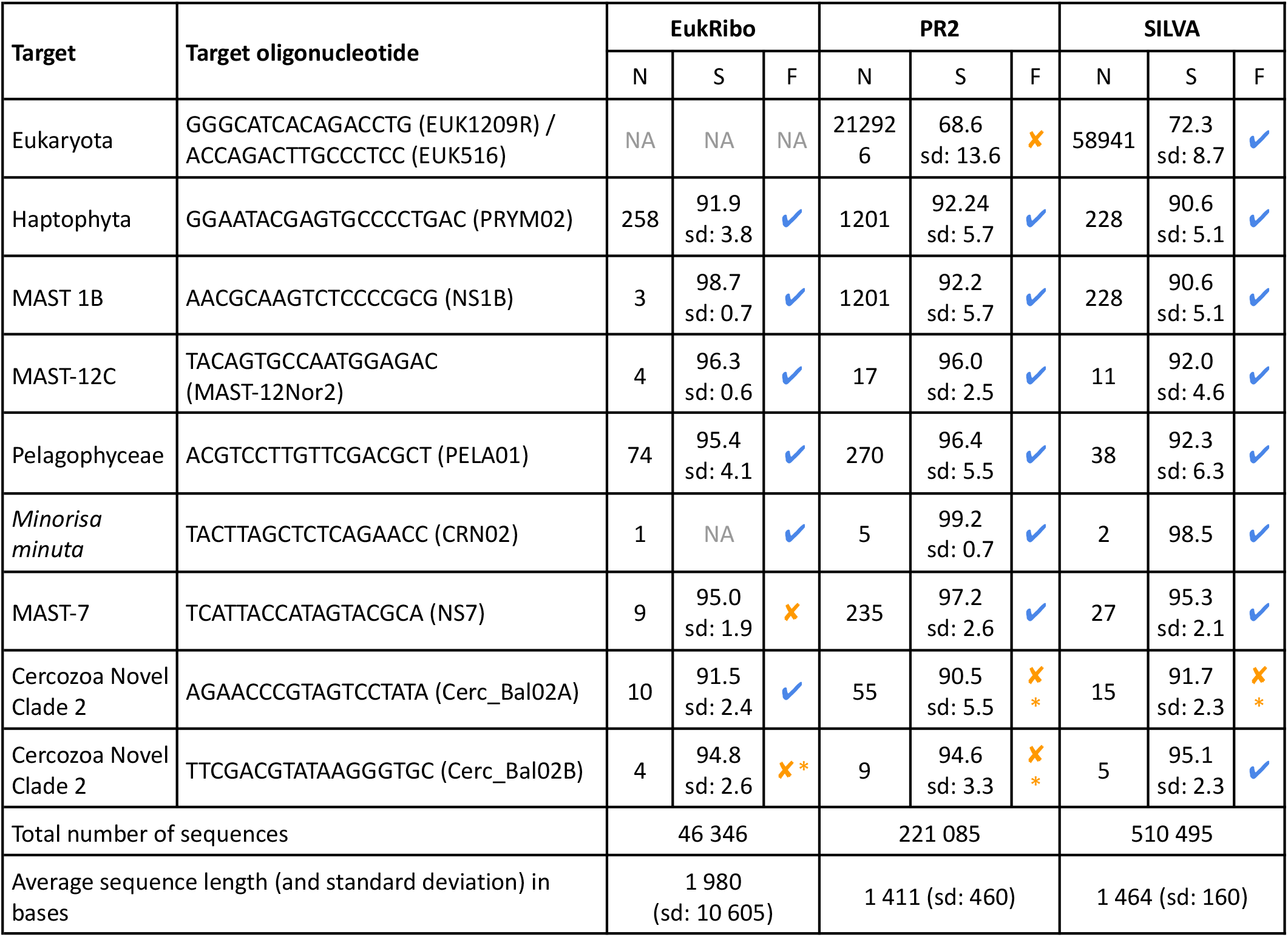
Qualitative performance of an unsupervised search for already published oligonucleotide probes using different databases (EukRibo, SILVA and PR2) showing the number of sequences in the target file (N), the average genetic diversity (measured as the average similarity identity by applying the --allpairs_global from VSEARCH) of the target file (S) and whether the given probe was found (✓) or not (✗) (F). *‘*NA’ represents ‘Not Applicable’ (i.e., lack of the target or excluding group in the given data), ‘sd’ represents ‘standard deviation’ of the given average, and ‘*’ after ‘✗’ means the probe was found after thorough manual curation of the target file.

General eukaryotic probes (such as EUK1209R; Giovannoni et al., 1988; or EUK516; Amann et al., 1990) were only retrieved in the SILVA database. It is known that general probes do not cover all targeted diversity (Vaulot et al., 2022), and these general probes were not found in PR2 mostly due to the extensive coverage of eukaryotic sequences (and small proportional coverage of prokaryotes) compared to the SILVA database. Both EUK1209R and EUK516 hit ∼60% of eukaryotic sequences in PR2. For such reasons, no general eukaryotic oligonucleotide was identified using the PR2 database.

When it comes to specific oligonucleotides the success rate was overall higher. For example the probes PRYM02 specific to Haptophyta (Lange et al., 1996), NS1B specific to MAST-1B (Massana et al., 2006), MAST-12Nor2 specific to MAST-12C; (Kolodziej and Stoeck, 2007) and PELA01 specific to Pelagophyceae (Simon et al., 2000) were identified in all 3 databases. The probe CRN02 specific to *Minorisa minuta* (Campo et al., 2013) was also found in all three databases, however these databases had to be curated independently and in detail since the three databases named *Minorisa minuta* sequences differently. While EukRibo and PR2 had only 1 and 5 sequences identified as *‘Minorisa minuta*’ respectively, SILVA had no sequences identified at the species level (but as ‘*Minorisa* sp.’) and therefore specific sequences had to be extracted by matching accession numbers.

The probe NS7 specific to MAST-7 (Giner et al., 2016) was only found in PR2 and SILVA, because it has a low specificity to clades MAST-7C and MAST-7D (which has 1 mismatch difference) and EukRibo contains only non-redundant diversity (i.e., the 163 sequences assigned to MAST-7A had a similarity of 99.1% (sd:0.7%) between them). Such selection of sequences reduces the 235 sequences assigned to MAST-7 from PR2 to just 9 in EukRibo, and thus the probe NS7 matches 66.6% of sequences in EukRibo, 87.7% in PR2 and 81.5% in SILVA. Lastly, this subclade diversity was the reason that the probes Cerc_Bal02A and Cerc_Bal02B, specific to different subclades within Cercozoa-Novel-Clade-2 (Piwosz and Pernthaler, 2011), were found in only one database each. Both Cerc_Bal02A and Cerc_Bal02B probes are specific to different sets of non-monophyletic diversity comprising around 37% and 11% of Cercozoa-Novel-Clade-2 sequences. Therefore, to retrieve such probes a manual selection of specific sequences according to the original scientific question is needed, in which case the probe is found.

Based on these observations, and further usage of the oligoN-design tools, we have included “Common problems and misconceptions” and “Good practices” sections in the manual documentation, where we have explained in detail recommendations and we will continuously update in the future. Briefly, we strongly recommend that, regardless of the software chosen for oligonucleotide design, to begin with a literature research and to further understand the diversity of your group of interest (by for example building a phylogenetic tree). When using oligoN-design, both the target group and input files should be properly examined and should contain the diversity that best matches the given scientific question. The oligoN-design tool will facilitate the design of specific oligonucleotides, although it is the user’s responsibility to provide a properly curated target and excluding files for the accurate detection of specific oligonucleotides.

Oligonucleotide design is a tedious work that requires a final empirical test for its completion. Therefore, bioinformatic pipelines will only provide theoretical candidate regions that have to be tested in the laboratory. Additional post-hoc resources can contribute to the *in silico* selection of experimental conditions for the selected oligonucleotides, such as oligoCalc and/or properties calculator. These resources are capable of estimating very accurately theoretical melting temperature and/or formamide concentration, and therefore increase the success during empirical tests in the wet lab.

## 5. Quantitative performance

The performance of the most computationally demanding functions was measured in time (Fig. 4), and in memory usage against both randomly generated fasta files and empirical datasets. Random fasta files were generated with the function fastaRandom.py, generating sequences of 1000 bases long. To recreate the different **target** and **excluding** files, the first sequence was randomly generated and the rest of the sequences contained a 1% mutation probability compared to the first sequence for the **target** file and 20% for the **excluding** file. Different file sizes were created containing from 10 up to 500,000 sequences (in a logarithmic series). Performance analyses were run on an Ubuntu 24.04 laptop with 32 GB RAM and 16 physical cores (Intel® Core™ i5-1250P).

**Figure 4.**
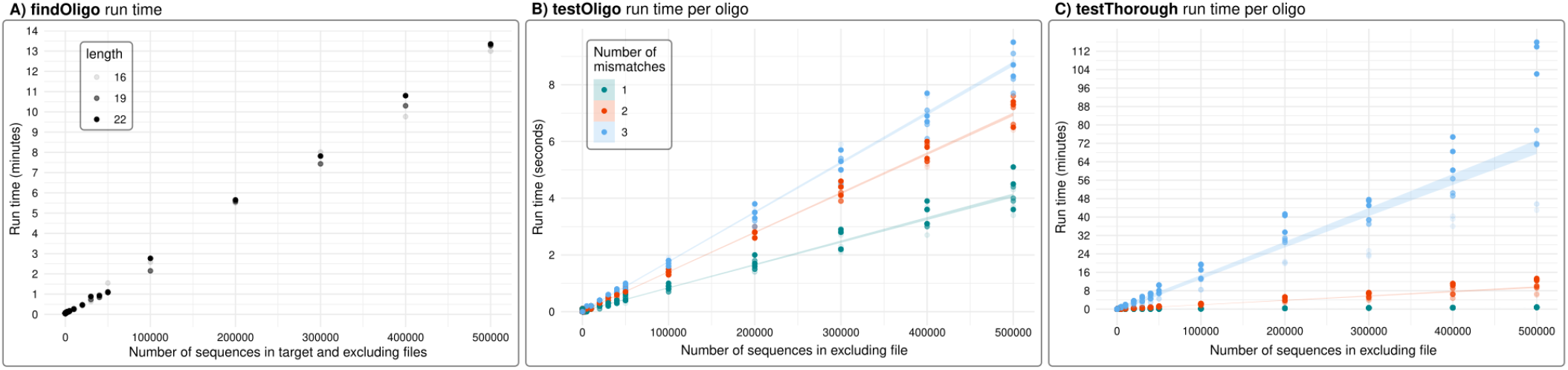
Run time of the three most computationally demanding functions using randomly-generated fasta files of different numbers of sequences (see text for further details), and comparing oligonucleotides of different lengths (16, 19 and 22 bases). **A)** Total run time in minutes of the function **findOligo** considering equal sizes for the **target** and **excluding** files. However, note that in most practical cases the **target** file will be significantly smaller than the excluding file, speeding up computation time. **B)** Run time per oligonucleotide in seconds of the function **testOligo** considering 1, 2 and 3 mismatches for different oligonucleotide lengths. **C)** Run time per oligonucleotide in minutes of the function **testThrough** considering 1, 2 and 3 mismatches for different oligonucleotide lengths.

When it comes to run time, **findOligo, testOligo** and especially **testThorough** are the most expensive functions. They all showed an increase in computing time that was directly proportional to the number of sequences in the fasta files (Fig. 4). The function **findOligo** showed a linear slope of 0.0015 seconds per sequence (of the **target** and **excluding** files) and no significant differences in the length of the oligonucleotide searched (Fig. 4A). When mismatches were tested, the number of mismatches more significantly increased the running time than the size of the fasta file. The function **testOligo** showed slopes of 0.0081, 0.014 and 0.017 seconds per oligo compared against 1000 sequences when allowing 1, 2 and 3 mismatches respectively (Fig. 4B). The slowest function of all is **testThorough**, showing slopes of 0.091, 1.14 and up to 8.48 seconds per oligo compared against 1000 sequences on average when allowing 1, 2 and 3 mismatches respectively (Fig. 4C). In addition, **testThorough** showed significant differences in the length of the oligonucleotide tested (linear regression p-value = 0.00494). Given these results, we recommend to use the **testOligo** function to curate the oligonucleotides and apply a thorough test only in those highly relevant oligonucleotides with the function **testThorough**.

The most expensive functions in terms of memory usage are **findOligo** and **getHomolog**, consuming up to ∼6 and ∼18 GB respectively when using the largest **target** and **excluding** files (with 500,000 sequences each; data not shown). Unlike run time, the memory usage of **findOligo** depends mostly on the number of k-mers in the given fasta file rather than the number of sequences, with up to 2 GB of memory difference in fasta files with different numbers of k-mers but similar number of sequences. While **findOligo** requires memory that is affordable in most modern desktop and laptop computers, **getHomolog** might require exceptional resources when considering very large datasets (>∼400k sequences). The rest of the functions use a similar memory as the file size, being up to 1GB in the largest examples.

## 6. Conclusion

With oligoN-design, we attempted to generate a simple and versatile tool for the design of specific oligonucleotides that can be integrated into bioinformatic pipelines, requiring only 2 fasta files as input. Based on our experience in primer and probe design with a wide diversity of different approaches, here we streamlined and automatized the complementary expertise of the authors to facilitate the development of specific probes and primers. We will continue to improve the accuracy, speed and robustness of oligoN-design in the future, adding new features and integrating further feedback from the community.

## 7. Acknowledgements

This work was supported by postdoctoral fellowships for MMS from the *Beatriu de Pinós* programme of the Government of Catalonia’s Secretariat for Universities and Research of the Generalitat de Catalunya Economy and Knowledge (grant no. 2021BP00068) and the European Union’s Horizon Europe research and innovation programme (grant no. 101103530; MSCA postdoctoral fellowship IMAGEN3D). MW was supported by a postdoctoral Benjamin Franklin Fellowship (project 464344344) from the German Research Council (Deutsche Forschungsgemeinschaft, DFG) and received funding from the European Union’s Horizon Europe Programme BlueRemediomics (grant agreement no. 101082304). This project has furthermore received funding from the European Research Council (ERC) under the European Union’s Horizon 2020 research and innovation programme (grant agreement no. 949745), from grants PID2023-152955NA-I00 and PID2022-137508NB-I00 funded by MICIU/AEI/10.13039/501100011033 and by ERDF/EU, and received support from the Departament de Recerca i Universitats de la Generalitat de Catalunya (exp. 2021 SGR 00751). We are grateful to the Roscoff Bioinformatics platform ABiMS (http://abims.sb-roscoff.fr), part of the Institut Français de Bioinformatique (ANR-11-INBS-0013) and BioGenouest network, for providing help and computing resources. We would like to thank Megan Sørensen, Charles Bachy and Anne Walraven for providing valuable feedback and testing previous versions of this tool.

## 8. Supplementary materials

**Figure S1.**
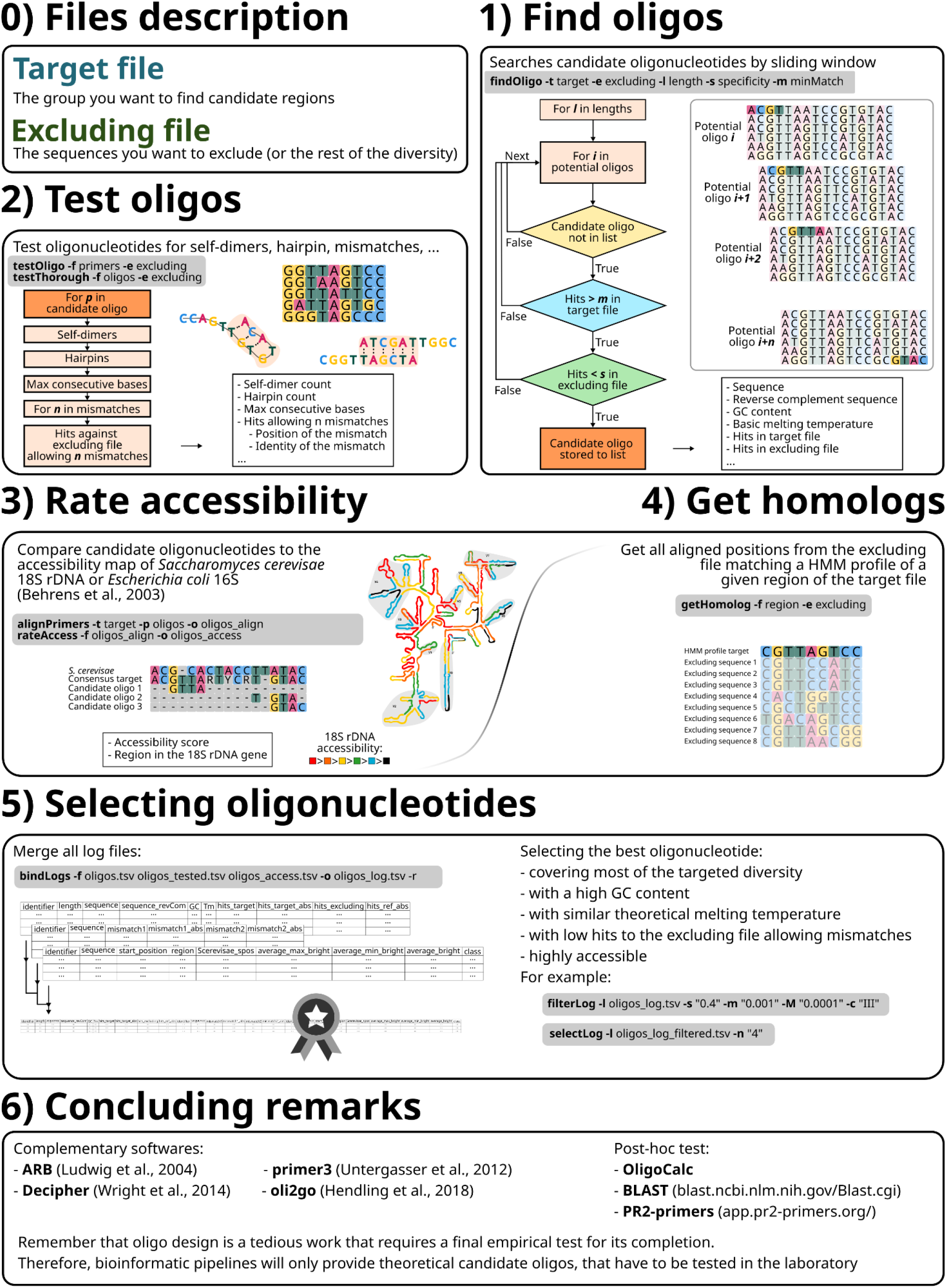
A detailed overview of the oligoN-design main functions.

