## Supplementary material for "OligoN-design: A simple and versatile tool to design specific probes and primers from large heterogeneous datasets": Figure S1

### 0) Files description

#### Target file
The group you want to find candidate regions

#### Excluding file
The sequences you want to exclude (or the rest of the diversity)

### 1) Find oligos

Searches candidate oligonucleotides by sliding window

**findOligo -t** target **-e** excluding **-l** length **-s** specificity **-m** minMatch

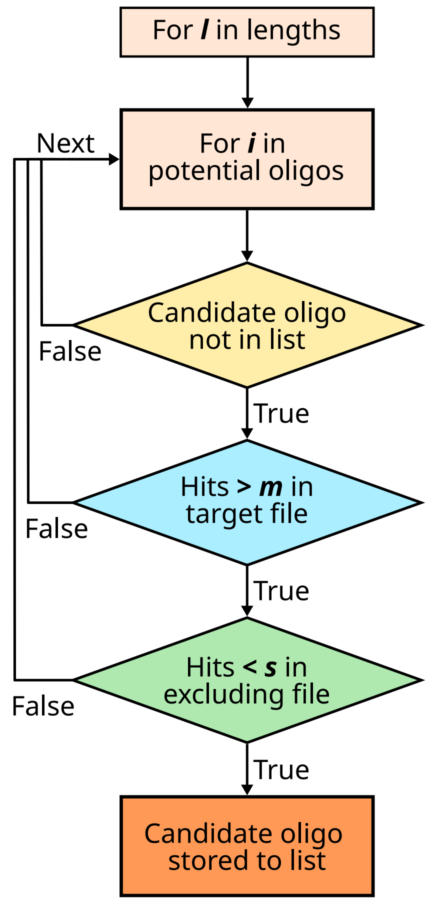


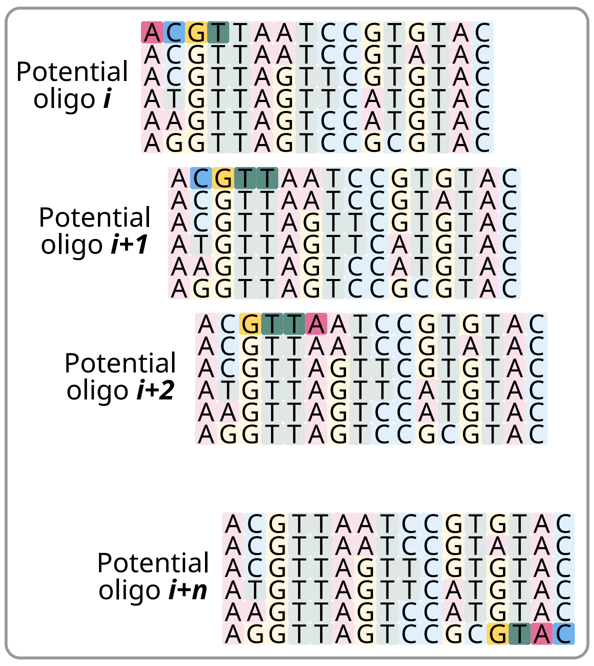


- Sequence
- Reverse complement sequence
- GC content
- Basic melting temperature
- Hits in target file
- Hits in excluding file
...

### 2) Test oligos

Test oligonucleotides for self-dimers, hairpin, mismatches, ...

**testOligo -f** primers **-e** excluding
**testThorough -f** oligos **-e** excluding

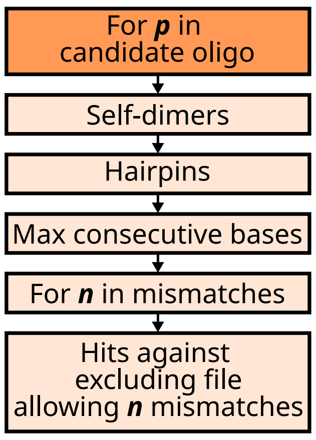


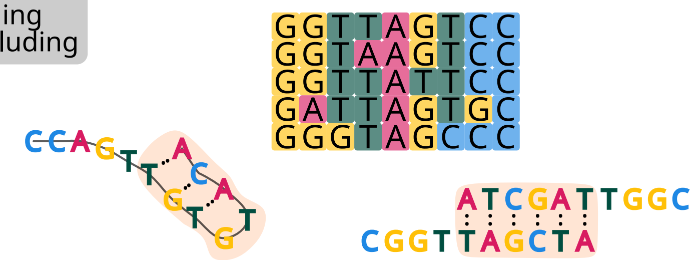

- Self-dimer count
- Hairpin count
- Max consecutive bases
- Hits allowing n mismatches
  - Position of the mismatch
  - Identity of the mismatch
...

### 3) Rate accessibility

Compare candidate oligonucleotides to the accessibility map of *Saccharomyces cerevisae* 18S rDNA or *Escherichia coli* 16S (Behrens et al., 2003)

**alignPrimers -t** target.fasta **-p** oligos.fasta **-o** oligos_align.fasta
**rateAccess -f oligos**_align.fasta **-o** oligos_access.tsv

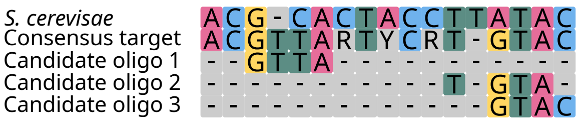


- Accessibility score
- Region in the 18S rDNA gene

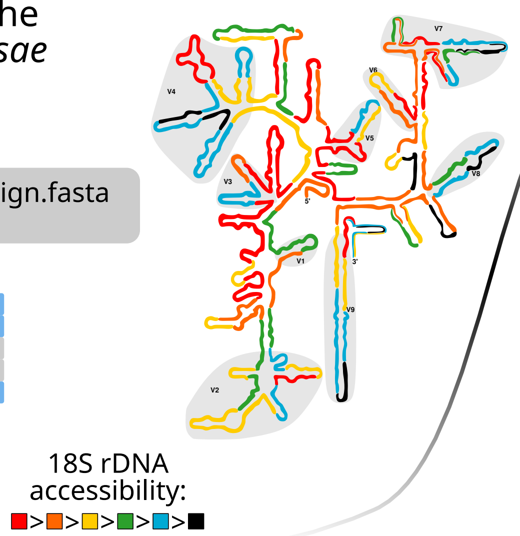


### 4) Get homologs

Get all aligned positions from the excluding file matching a HMM profile of a given region of the target file

**getHomolog -f** region.fasta **-e** excluding.fasta

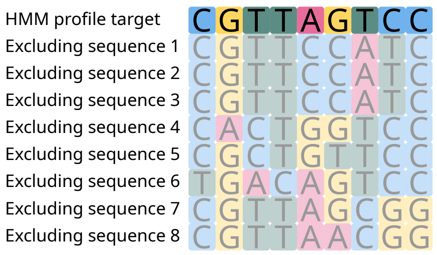


### 5) Selecting oligonucleotides

Merge all log files:

**bindLogs -f** oligos.tsv oligos_tested.tsv oligos_access.tsv **-o** oligos_log.tsv -r

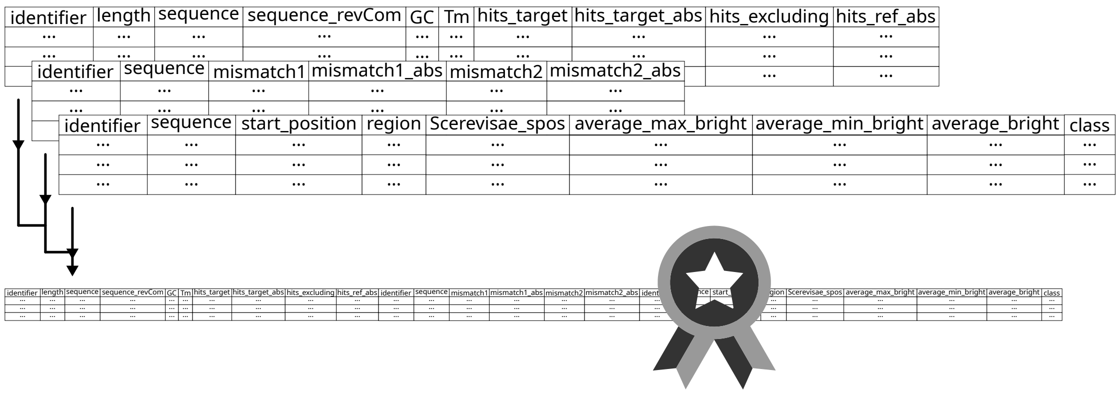


Selecting the best oligonucleotide:
- covering most of the targeted diversity
- with a high GC content
- with similar theoretical melting temperature
- with low hits to the excluding file allowing mismatches
- highly accessible

For example:

**filterLog -l** oligos_log.tsv **-s** "0.4" **-m** "0.001" **-M** "0.0001" **-c** "III"

**selectLog -l** oligos_log_filtered.tsv **-n** "4"

### 6) Concluding remarks

Complementary softwares:
- **ARB** (Ludwig et al., 2004)
- **Decipher** (Wright et al., 2014)
- **primer3** (Untergasser et al., 2012)
- **oli2go** (Hendling et al., 2018)

Post-hoc test:
- **OligoCalc**
- **BLAST** (blast.ncbi.nlm.nih.gov/Blast.cgi)
- **PR2-primers** (app.pr2-primers.org/)

Remember that oligo design is a tedious work that requires a final empirical test for its completion.
Therefore, bioinformatic pipelines will only provide theoretical candidate oligos, that have to be tested in the laboratory
